# A novel conformational state for SARS-CoV-2 main protease

**DOI:** 10.1101/2021.03.04.433882

**Authors:** Emanuele Fornasier, Maria Ludovica Macchia, Gabriele Giachin, Alice Sosic, Matteo Pavan, Mattia Sturlese, Cristiano Salata, Stefano Moro, Barbara Gatto, Massimo Bellanda, Roberto Battistutta

## Abstract

The SARS-CoV-2 main protease (M_pro_) has a pivotal role in mediating viral genome replication and transcription of coronavirus, making it a promising target for drugs against Covid-19 pandemic. Here we present a crystal structure of M_pro_ disclosing new structural features of key regions of the enzyme. We show that the oxyanion loop, involved in substrate recognition and enzymatic activity, can adopt a new conformation, which is stable and significantly different from the known ones. In this new state the S1 subsite of the substrate binding region is completely reshaped and a new cavity near the S2’ subsite is created. This new structural information expands the knowledge of the conformational space available to M_pro_, paving the way for the design of novel classes of inhibitors specifically designed to target this unprecedented binding site conformation, thus enlarging the chemical space for urgent antiviral drugs against Covid-19 pandemic.

## Introduction

M_pro_ is a cysteine peptidase essential for the replication of SARS-CoV-2 (Wu et al., 2020), with 96% sequence identity and very similar 3D structure to SARS-CoV M_pro_ (Anand et al., 2003; Douangamath et al., 2020; Jin et al., 2020a, 2020b; Kneller et al., 2020c; Yang et al., 2003; Zhang et al., 2020). M_pro_ is involved in the proteolytic processing of the large polyproteins pp1a and pp1ab with the formation of individual non-structural proteins (Snijder et al., 2016). M_pro_ forms a homodimer fundamental for the proper catalytic activity (Anand et al., 2002). A key role is played by the “N-finger” as the N-terminal tail of one protomer interacts and stabilizes the binding site (S1 subsite) of the other protomer (Verschueren et al., 2008). The N-finger and the C-terminus are the result of the autoproteolytic processing of M_pro_. In mature enzyme both termini of one protomer are facing the active side of the other. The enzymatic cleavage of the substrate occurs at the C-terminal end of a conserved glutamine in position P1 of the consensus sequence, with His41 and Cys145 as catalytic dyad (Anand et al., 2002). An important structural element for the catalytic event is the oxyanion loop, residues 138-145, lining the binding site for glutamine P1 and assumed involved in the stabilization of the tetrahedral acyl transition state (Anand et al., 2002; Lee et al., 2020; Verschueren et al., 2008). In the vast majority of structures, the oxyanion loop adopts the same conformation (here called “canonical”) (Douangamath et al., 2020; Jin et al., 2020a, 2020b; Zhang et al., 2020), whose mobility has been described by room temperature x-ray crystallography (Kneller et al., 2020b, 2020a). In few cases it can also exist in a different “collapsed” conformation, considered catalytically incompetent, often associated with modifications in the N- or C-termini or with acidic pH values (Verschueren et al., 2008; Yang et al., 2003).

## Results and Discussion

In a campaign to get structural insights on SARS-CoV-2 M_pro_ we analyzed 27 different datasets to determine the crystal structure of M_pro_ in complex with inhibitors, namely masitinib, manidipine and bedaquiline (Ghahremanpour et al., 2020). Among these, as “positive” controls (i.e. structures already known) there were ligand-free M_pro_ and M_pro_ in complex with inhibitor boceprevir (Fu et al., 2020). Almost all tested crystals were monoclinic (space group C2, Table SI), isomorphous to the crystals of the free enzyme 6Y2E (Zhang et al., 2020) and to most of the deposited M_pro_ structures, indicating the same crystal contacts. After successful MR and a first round of refinement, in most cases the electron density was clearly visible along the entire sequence, indicating a protein matrix with a structure very similar to the search models 6Y2E and 5REL (including the complex with boceprevir). However, there were a significant number of cases, around 10, where the electron density was of much lower quality or even absent in particular portions of the protein, namely for residues 139-144 of the oxyanion loop, residues 1-3 of the N-finger and the side chain of His163 in the S1 specificity subsite, all residues part of the active site. To cope with the known molecular replacement bias problem and to correctly rebuild the ambiguous parts, we performed new MR runs using as search model structure 6Y2E deprived of residues 139-144 and 1-3, and with an alanine instead of a histidine at position 163 (to remove the His side chain). This allowed to confirm perturbations in the conformation of the selected areas for 10 structures while clear electron densities were visible for the remaining cases with the oxyanion loop unambiguously in the canonical conformation (Fig. S1 and S2).

In some cases, the electron density was so poor that the tracing of the chain was very problematic, and it was not possible to reliably rebuild entirely the mobile zones (Fig. S1b). For four structures it was possible to efficiently model residues 139-144, 1-3 and the side chain of His163 in “new” conformations, different from the “canonical” and “collapsed” ones (Fig. S1c). In summary, we found three different conformational states for the oxyanion loop: canonical (Fig. S1a), flexible (i.e. with poor electron density, Fig. S1b) and, strikingly, in a new state (Fig. S1c), clearly different from the canonical and the collapsed ones (Fig. S3). All structures, including the one with the oxyanion loop in the new state (hereinafter called “new-M_pro_”), refer to a correctly autoprocessed and functional protein, produced and crystallized with procedures similar to those of canonical 6Y2E (Zhang et al., 2020), showing the same crystal contacts (see Methods). All structures are the result of very similar experimental procedures, performing multi-parallel experiments, with no apparent reasons for this diversity of conformations if not for the coexistence in solution of different conformational states (denoting high mobility) for the oxyanion loop, in mutual equilibrium.

Here we describe one of the structures of M_pro_ determined in the new state (no relevant differences exist between the four new-M_pro_ structures). Data collection and final model statistics are reported in Table SI, final electron densities for the oxyanion region in Fig. 1.

**Fig. 1.**
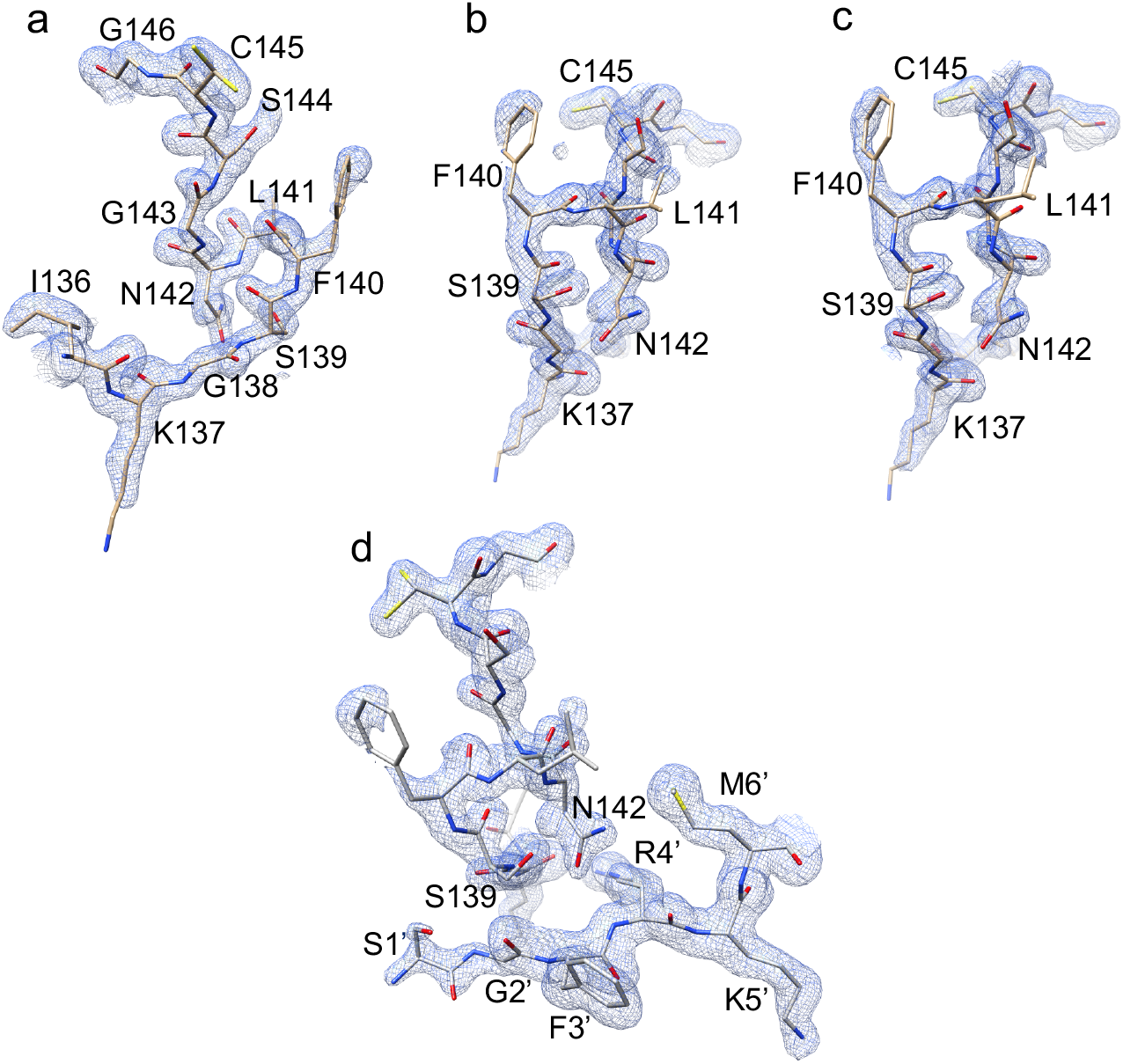
Final electron densities. 2F_o_-F_C_ maps contoured at 1.0 σ level are shown. In panel a and b two views of the final electron density for the oxyanion loop in the new conformation. Leu141 and solvent exposed Phe140 and Lys137 side chains have incomplete densities indicating various degree of flexibility. Panel c, simulated-annealing omit map (oxyanion loop residues 138-146 were omitted), view as in b. Panel d, electron density in the inter protomers (intra-dimer) interaction area between the oxyanion loop of one protomer and the N-finger of the other protomer (residues Ser1’-Met6’).

The most striking property of new-M_pro_ is the different conformational state of the oxyanion loop characterized by two consecutive ß-turns with hydrogen bonds between Ser139-CO and Gln142-NH and between Leu141-CO and Ser144-NH (Fig. 2a). The loop is stabilized by other hydrogen bonds. In the new conformation Asn142-Ca and the side-chain of Phe140 move away from the canonical position of 9.8 Å and around 7.5 Å, respectively (Fig. 2b). Notably, Gly143-NH, assumed stabilizing the tetrahedral oxyanion intermediate during catalysis (Anand et al., 2002; Lee et al., 2020; Verschueren et al., 2008), is moved 8.8 Å apart, raising the possibility that this new conformation is catalytically incompetent. However, the position of the catalytic dyad is not altered (Fig. S4), with Cys145 side chain in double conformation.

**Fig. 2.**
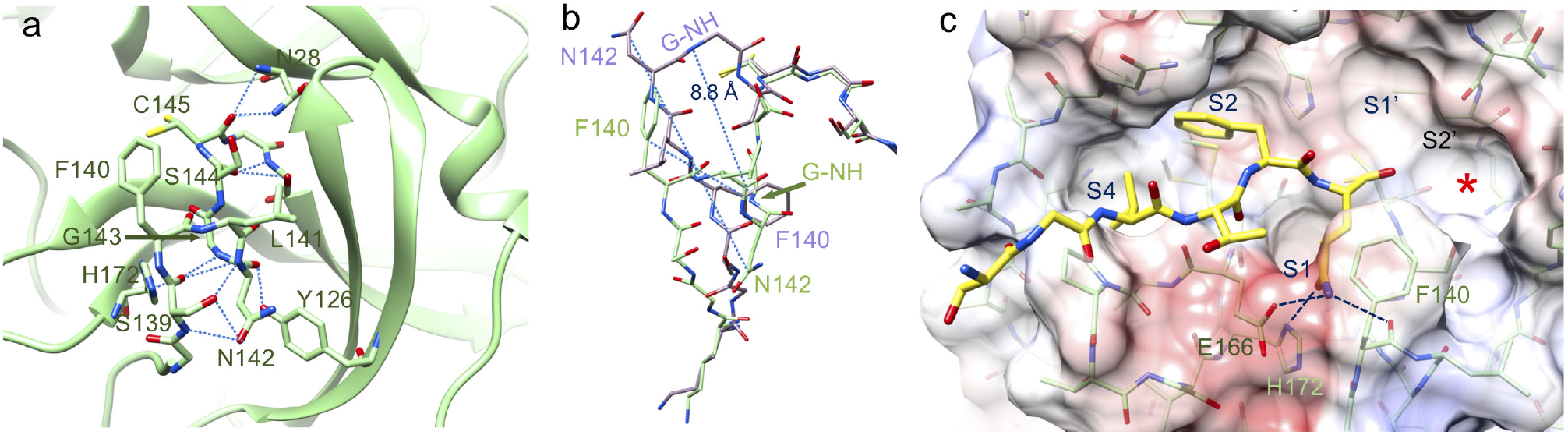
Details of new-M_pro_. a, the new conformation of the oxyanion loop is stabilized by several hydrogen bonds (blue dashed lines). The side-chain of catalytic Cys145 has a double conformation. b, comparison between new (green) and canonical (light magenta, PDB-ID 6Y2E) oxyanion loop. There are large movements (blue dashed lines) for the side-chains of Asn142 and Phe140. Gly143-NH (G-NH), involved in the stabilization of the tetrahedral intermediate, moves 8.8 Å away. c, substrate glutamine in S1 subsite of new-M_pro_ (modelled based on the acyl-intermediate 7KHP, in yellow) can still interact with Glu166-Oε and Phe140-CO, even if with a different geometry. His163 is no more available for binding but can be replaced by His172 that moves towards the S1 subsite. The new cavity near the S2’ subsite is indicated by a red asterisk.

The superposition of the new conformation with the canonical one in the complex with the acyl-intermediate 7KHP (Lee et al., 2020) does not show evident steric clashes for the substrate, indicating that new-M_pro_ could bind P1 glutamine with Glu166-Oε and Phe140-CO, even if with a different geometry (Fig. 2c). Although His163 is no more available for binding as it rotates away to avoid steric clashes with Gly143-CO (Fig. 3a), it can be replaced by His172 that moves towards the S1 subsite (Fig. 3a and 2c). As consequence of the new oxyanion conformation of one protomer, residues 1-3 of the N-finger of the other protomer (protomer’) moves away (Fig. 3b and 3c), with Gly2’-CO at 3.2 Å from Ser139-NH. Remarkably, Arg4’ does not move and the inter-protomers salt bridge with Glu290 (important for catalytic activity and dimer formation) is still present (Fig. 3b). Another characteristic of the new structure is the destabilization of the C-terminal tail whose electron density is not visible anymore from residue 301 on, indicating high flexibility. This is due to the rearrangement of the interactions between the oxyanion loop of one protomer and the N-finger and C-terminal portion of the other (Fig. 4).

**Fig. 3.**
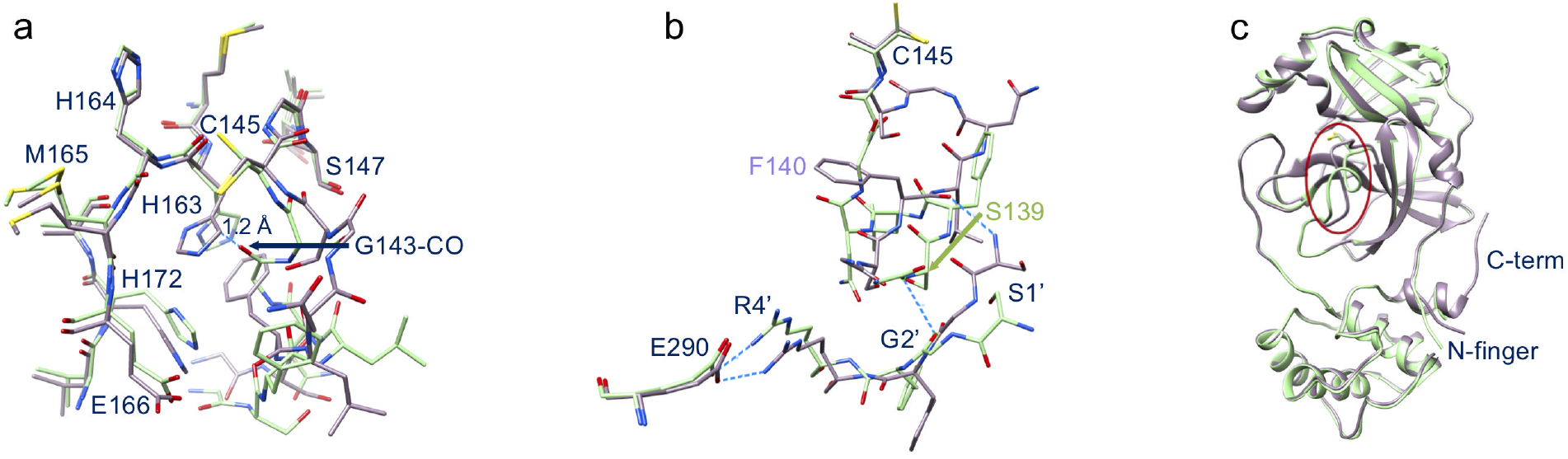
Comparison between new (green) and canonical (light magenta) M_pro_. a, in the new structure the side chain of His163 rotates away to avoid steric clashes with the oxyanion loop: in the canonical conformation (PDB-ID 6Y2E) the His163 side chain is at 1.2 Å from the new position of Gly143-CO. Note also the movement of His172. b, the new oxyanion loop of one protomer pushes away residues 1’-3’ of the other protomer; however, the key salt-bridge between Arg4’ and Glu290 is conserved. c, overall superposition of canonical and new-M_pro_ shows that, besides in the oxyanion loop (red ellipsoid), major differences are located in the N-finger and the C-terminal tail (not visible in new-M_pro_).

**Fig. 4.**
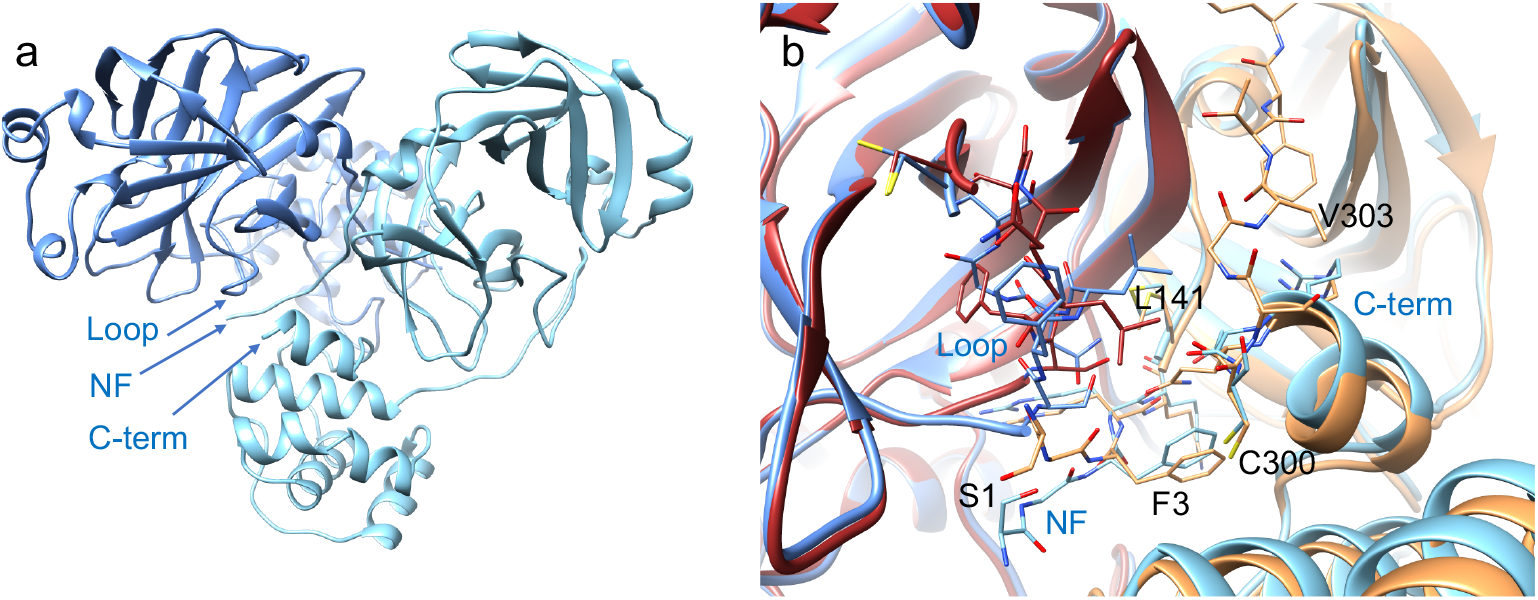
Dimeric architecture of M_pro_. a, the new conformation of the oxyanion loop (loop) causes changes in the interface between protomer-A (blue) and protomer-B (light blue) at the level of the N-finger (NF) and the C-terminal tail (C-term). b, local differences between the new structure (blue-based colors as in a) and canonical 6Y2E (brown-based colors, with intact C-terminus): shift of Leu141 side chain seems to have major effects in destabilizing the C-terminal tail of the new structure.

Notably, the new conformation of the oxyanion loop generates a new cavity near position S2’ as evident from the comparison of the new structure and the SARS-CoV-2 acyl-enzyme 7KHP (Lee et al., 2020) and the SARS-CoV 11_mer_ substrate complex 2Q6G (Xue et al., 2008) (Fig. S5 and Fig 2c).

This new structure was derived from crystals obtained with M_pro_ pre-incubated with inhibitors masitinib, manidipine or bedaquiline, however in no cases electron densities indicating the presence of the inhibitors were detected. This is explainable by the reported medium/low IC_50_ (in the range 2.5-19 μM) (Drayman et al., 2020; Ghahremanpour et al., 2020) and by the very low aqueous solubility of the molecules (when inhibitors in 100% DMSO were added to the protein solution visible white precipitates appeared). Notably, some structures from crystals coming from co-crystallization experiments with the same inhibitors pre-incubated with M_pro_ show the oxyanion canonical conformation, indicating that these molecules are not strict determinants for the new conformation. However, it seems that these inhibitors can somehow favor the new conformation of the oxyanion loop, which is enough stable to be selected by the crystallization process.

To test the stability and to model dynamics of new-M_pro_, in particular of the oxyanion loop, we performed crystallographic ensemble refinement (Burnley et al., 2012) and MD simulations. The 60 structures generated by ensemble refinement, compatible with the crystallographic restraints, confirm the new conformation of the oxyanion loop, and reveal that its flexibility is not higher than that of other portions of the substrate-binding region (residues 43-51 and 188-198) (Fig. S6). Interestingly, in 4 out of 60 structures the oxyanion loop conformation is similar to the canonical one, in line with the observation of a residual electron density compatible with the presence of a small fraction of this conformation.

1 μs MD simulations show that both the canonical and the new structure maintain the initial state, denoting the stability of the new oxyanion loop conformation (Fig. S7). A pronounced fluctuation of the C-terminus of new-M_pro_ is observed in agreement with the absence of electronic density for this region. The mobility pattern displayed by MD matches that deduced by the ensemble refinement approach, in particular for regions 43-51 and 188-198, part of the substrate binding region (S3 and S4 subsites), and region 272-279 in the C-terminal domain III (Fig. S6 and S7). In summary, we had the opportunity to capture a new state for M_pro_ expanding the knowledge of the conformational space accessible to the enzyme. The remodeling of the S1 subsite in new-M_pro_ and the formation of the nearby cavity offer attractive possibilities for the design of novel antiviral drugs targeting M_pro_. The plasticity properties of new-M_pro_ are also relevant for the debated issue whether M_pro_ proteolytic recognition is based on structural selection or on substrate-induced subsite cooperativity (Behnam, 2021). Intriguing is also the possibility that the remodeling of the S2’ subsite can be correlated to the high amino acid variation in position P2’ of SARS coronaviruses non-structural proteins (nsp) cleavage sites, M_pro_ autoprocessing included (Behnam, 2021).

## METHODS

### Protein expression and purification

The plasmid PGEX-6p-1 encoding SARS-CoV-2 M_pro_ (Zhang et al., 2020) was a generous gift of Prof. Rolf Hilgenfeld, University of Lübeck, Lübeck, Germany. Protein expression and purification were adapted from Zhang, L. *et al.* (Zhang et al., 2020) (where the structure of M_pro_ 6Y2E in the canonical form was presented). The expression plasmid was transformed into *E. coli* strain BL21 (DE3) and then pre-cultured in YT medium at 37 °C (100 μg/mL ampicillin) overnight. The preculture was used to inoculate fresh YT medium supplied with antibiotic and the cells were grown at 37 °C to an OD600 of 0.6–0.8 before induction of overexpression with 0.5 mM isopropyl-D-thiogalactoside (IPTG). After 5 h at 37 °C cells were harvested by centrifugation (5000g, 4 °C, 15 min) and frozen. The pellets were resuspended in buffer A (20 mM Tris, 150 mM NaCl, pH 7.8) supplemented with lysozyme, DNase I and PMSF for the lysis. The lysate was clarified by centrifugation at 12000 g at 4 °C for 1 h and loaded onto a HisTrap HP column (GE Healthcare) equilibrated with 98% buffer A/2% buffer B (20 mM Tris, 150 mM NaCl, 500 mM imidazole, pH 7.8). The column was washed with 95% buffer A/5% buffer B and then His-tagged M_pro_ was eluted with a linear gradient of imidazole ranging from 25 mM to 500 mM. Pooled fractions containing target protein was subjected to buffer exchange with buffer A using a HiPrep 26/10 desalting column (GE Healthcare). Next, PreScission protease was added to remove the C-terminal His tag (20 μg of PreScission protease per mg of target protein) at 12 °C overnight. Protein solution was loaded onto a HisTrap HP column connected to a GSTtrap FF column (GE Healthcare) equilibrated in buffer A to remove the GST-tagged PreScission protease, the His-tag, and the uncleaved protein. M_pro_ was finally purified with a Superdex 75 prep-grade 16/60 (GE Healthcare) SEC column equilibrated with buffer C (20 mM Tris, 150 mM NaCl, 1 mM EDTA, 1 mM DTT, pH 7.8). Fractions containing the target protein at high purity were pooled, concentrated at 25 mg/ml and flash-frozen in liquid nitrogen for storage in small aliquots at −80 °C.

### Protein characterization and enzymatic kinetics

Correctness of M_pro_ DNA sequence was verified by sequencing the expression plasmid. The molecular mass was determined as follows: the recombinant SARS-CoV-2 M_pro_, diluted in 50% acetonitrile with 0.1% of formic acid, was analyzed by direct infusion electrospray ionization (ESI) on a Xevo G2-XS QTOF mass spectrometer (Waters). The detected species displayed a mass of 33796.64 Da, which matches very closely the value of 33796.81 Da calculated from the theoretical full-length protein sequence (residues 1-306). A representative ESI-MS spectrum is reported in Fig. S8. To characterize the enzymatic activity of our recombinant M_pro_, we adopted a FRET-based assay using the substrate 5-FAM-AVLQISGFRK(DABCYL)K (Proteogenix). The assay was performed by mixing 0.05 μM M_pro_ with different concentrations of substrate (1-128 μM) in a buffer composed of 20 mM Tris, 100 mM NaCl, 1 mM EDTA, 1 mM DTT, pH 7.3. Fluorescence intensity (Ex = 485 nm/Em = 535 nm) was monitored at 37 °C with a microplate reader VictorIII (Perkin Elmer). A calibration curve was created by measuring multiple concentrations (from 0.001 to 5 μM) of free fluorescein in a final volume of 100 μL reaction buffer. Initial velocities were determined from the linear section of the curve, and the corresponding relative fluorescence units per unit of time (ΔRFU/s) was converted to the amount of the cleaved substrate per unit of time (μM/s) by fitting to the calibration curve of free fluorescein. The catalytic efficiency kcat/Km resulted 4819± 399 s^−1^ M^−1^, in line with literature data (Ma et al., 2020; Zhang et al., 2020).

### Crystallization and data collection

A frozen aliquot of M_pro_ was thawed in ice, diluted in a 1:2 ratio with buffer C (20 mM Tris, 150 mM NaCl, 1 mM EDTA, 1 mM DTT, pH 7.8) to a final concentration of 12.5 mg/mL and cleared by centrifugation at 16000 *g.* Inhibitors masitinib, manidipine, bedaquiline and boceprevir were dissolved in 100% DMSO to a concentration of 100 mM. The protein was crystallized both in apo form and in presence of inhibitors by co-crystallization. In all cases, final crystal growth was obtained by microseeding starting from small crystals of the free enzyme. The protein in the apo form was crystallized using the sitting-drop vapor-diffusion method at 18 °C, mixing 1.0 μL of M_pro_ solution with 1.0 μL of precipitant solution (0.1 M MMT (DL-malic acid, MES, and Tris base in molar ratio 1:2:2), pH 7.0, 25% PEG 1500) with 0.2 μL of seed stock (diluted 1:500, 1:1000 or 1:2000 with precipitant solution) and equilibrated against a 300 μL reservoir of precipitant solution. Crystals appeared overnight and finished growing in less than 48 h after the crystallization drops were prepared. In the case of co-crystallization, M_pro_ was incubated for 16 h at 8 °C with 13-fold molar excess of inhibitor (final DMSO concentration 5%). After incubation a white precipitate appeared and the solutions were cleared by centrifugation at 16000 x g; then the protein was crystallized under the same conditions described for the apo form. For data collections, crystals were fished from the drops, cryo-protected with a quick deep into 30% PEG 400 (with 5 mM inhibitor in the case of co-crystals) and flash-cooled in liquid nitrogen. Crystals were monoclinic (space group C2, Table SI), isomorphous to the crystals of the free enzyme 6Y2E, with one monomer in the asymmetric unit, the functional dimer being formed by the crystallographic twofold axis.

### Structure determination and refinement

Data collections were performed at ESRF, beamlines ID23-2 and ID23-1. Diffraction data integration and scaling were performed with XDS (Kabsch, 2010), data reduction and analysis with Aimless (Evans and Murshudov, 2013). Initially, structures were solved by Molecular Replacement (MR) with Phaser (McCoy et al., 2007) from Phenix (Liebschner et al., 2019), using as search model structures 6Y2E and 5REL (M_pro_ in complex with PCM-0102340) (Douangamath et al., 2020). To limit MR model bias in critical zones (namely residues 139-144, 1-3 and the side chain of His163) we then performed new MR runs using as search model structure 6Y2E without residues 139-144 and 1-3, and with an alanine instead of an histidine at position 163. Only for cocrystallization experiments with boceprevir electron density relative to the ligand was clearly visible since the beginning of the refinement (Fig. S2), and the 3 final structures, modelled from residue 1 to 306 (to compare with the “new” structure modelled until residue 301), are virtually identical to the PDB deposited ones (Fu et al., 2020). In all other cases, no electron densities indicating the presence of inhibitors masitinib, manidipine or bedaquiline in the active site (or elsewhere) were detectable. For four structures it was possible to efficiently model residues 139-144, 1-3 and the side chain of His163 in “new” conformations. The final structures were obtained by alternating cycles of manual refinement with Coot (Emsley and Cowtan, 2004) and automatic refinement with phenix.refine (Afonine et al., 2012). The final electron density for the new oxyanion conformation and the N-finger are reported in Fig. 1. At the end, the model was submitted to ensamble.refinement (Burnley et al., 2012) by Phenix with default parameters. Statistics on data collection and refinement are reported in Table SI.

### Molecular Modeling

Molecular Dynamics trajectories were collected on a heterogeneous NVIDIA GPU cluster composed of 20 GPUs whose model span from GTX1080 to RTX2080Ti. For structures preparation, the coordinates of the canonical conformation of SARS-CoV-2 Main Protease (M_pro_) were retrieved from the Protein Data Bank (PDB-ID: 7K3T). Coordinates for both the canonical and the newly identified conformation of SARS-CoV-2 Main Protease were processed with the aid of Molecular Operating Environment (MOE) 2019.011 (“Chemical Computing Group (CCG) | Research,” n.d.) structure preparation tool. At first, the functional unit of the protease (the dimeric form) was restored applying a symmetric crystallographic transformation to each asymmetric unit. Residues with alternate conformation were assigned to the highest occupancy alternative. The last 6 residues of the non-canonical structures were added using MOE Loop Modeler tool. MOE Protonate3D tool was used to assign the most probable protonation state of each residue (pH 7.4, T = 310 K, i.f = 0.154). Finally, ions and each co-crystallized molecule except for water were removed. The system setup for the MD simulations was carried out using tleap software implemented in the AmberTools14 (Case et al., 2005) suite. AMBER ff14SB (Maier et al., 2015) was adopted for system parametrization and partial charges attribution. Protein structures were explicitly solvated in a rectangular prism TIP3P (Jorgensen et al., 1983) periodic water box whose borders were placed at a distance of 15 Å from any protein atoms. Na^+^ and Cl^−^ atoms were added to neutralize the system until a salt concentration of 0.154 M was reached. Molecular Dynamics simulations were then performed using ACEMD3 (Harvey et al., 2009) software, which is based upon OpenMM 7.4.2 (Eastman et al., 2017) engine. At first, 1000 steps of energy minimization were executed using the conjugate-gradient algorithm. Then, a two steps equilibration procedure was carried out: the first step consisted of 1 ns of canonical ensemble (NVT) simulation with 5 kcal mol^−1^A^−2^ harmonic positional constraints applied to each protein atom while the second one consisted of 1 ns of isothermal-isobaric (NPT) simulation with 5 kcal mol^−1^A^−2^ harmonic positional constraints applied only to protein alpha carbons. The production phase consisted of three independent MD replica for each protein conformation. Each simulation had a duration of 1 μs and was performed using the NVT ensemble at a constant temperature of 310 K with a timestep of 2 fs. For both the equilibration and the production stage, the temperature was maintained constant by a Langevin thermostat. During the second step of the equilibration stage, the pressure was maintained at the fixed value of 1 bar. For Molecular Dynamics simulation analysis, MD trajectories were aligned using protein α-carbon atoms from the first trajectory frame as a reference, wrapped into an image of the system under periodic boundary conditions (PBC) and subsequently saved using a 200 ps interval between each frame and removing any ion and water molecule using Visual Molecular Dynamics 1.9.2 (Humphrey et al., 1996) (VMD) software. Root Mean Squared Deviation (RMSD) and Root Mean Squared Fluctuation (RMSF) of atomic positions along the trajectory were calculated for protein alpha carbons exploiting the ProDy (Bakan et al., 2011) Python module. The collected data was then plotted making use of the Matplotlib (Hunter, 2007) Python library.

## Data availability

Coordinates and structure factors for SARS-CoV-2 new-M_pro_ have been deposited in the Protein Data Bank (PDB) under accession code 7NIJ.

## Acknowledgments

The authors are grateful to Prof. Rolf Hilgenfeld, University of Lübeck, Lübeck, Germany, for the free availability of plasmid encoding SARS-CoV-2 M_pro_ (Zhang et al., 2020). We thank ESRF beamline ID23-2 local contact Daniele De Sanctis and beamline ID23-1 local contact Romain Talon for help and assistance with data collection. This work was supported by funding from the CARIPARO Foundation (“Progetti di ricerca sul Covid-19” N. ***55812*** to BG and PhD-scholarship to EF), from the Department of Chemical Sciences (project P-DiSC #01BIRD2018-UNIPD to M.B.), from the University of Padua (Starting Grant STARS@UNIPD – call 2019 to GG).

## Author contributions

Protein expression and purification were planned and performed by MLM and MB, mass spectrometry by AS and BG, activity measurements by AS, MLM and BG, crystallization experiments and data collection by EF, GG and RB, data processing, structure solution and refinement by RB, MD simulation by MP, MS, and SM, overall design of the research by MB, BG, SM, CS and RB, manuscript writing by RB.

## Additional information

Supplementary information is available for this paper.

## References

Afonine PV, Grosse-Kunstleve RW, Echols N, Headd JJ, Moriarty NW, Mustyakimov M, Terwilliger TC, Urzhumtsev A, Zwart PH, Adams PD. 2012. Towards automated crystallographic structure refinement with phenix.refine. Acta Crystallogr D Biol Crystallogr 68:352–367. doi:10.1107/S0907444912001308

Anand K, Palm GJ, Mesters JR, Siddell SG, Ziebuhr J, Hilgenfeld R. 2002. Structure of coronavirus main proteinase reveals combination of a chymotrypsin fold with an extra alpha-helical domain. EMBO J 21:3213–3224. doi: 10.1093/emboj/cdf327

Anand K, Ziebuhr J, Wadhwani P, Mesters JR, Hilgenfeld R. 2003. Coronavirus main proteinase (3CLpro) structure: basis for design of anti-SARS drugs. Science 300:1763–1767. doi:10.1126/science.1085658

Bakan A, Meireles LM, Bahar I. 2011. ProDy: protein dynamics inferred from theory and experiments. Bioinforma Oxf Engl 27:1575–1577. doi:10.1093/bioinformatics/btr168

Behnam MAM. 2021. Protein structural heterogeneity: A hypothesis for the basis of proteolytic recognition by the main protease of SARS-CoV and SARS-CoV-2. Biochimie 182:177–184. doi:10.1016/j.biochi.2021.01.010

Burnley BT, Afonine PV, Adams PD, Gros P. 2012. Modelling dynamics in protein crystal structures by ensemble refinement. eLife 1:e00311. doi:10.7554/eLife.00311

Case DA, Cheatham TE, Darden T, Gohlke H, Luo R, Merz KM, Onufriev A, Simmerling C, Wang B, Woods RJ. 2005. The Amber biomolecular simulation programs. J Comput Chem 26:1668–1688. doi:10.1002/jcc.20290

Chemical Computing Group (CCG) | Research. n.d. https://www.chemcomp.com/Research.htm#Citations

Douangamath A, Fearon D, Gehrtz P, Krojer T, Lukacik P, Owen CD, Resnick E, Strain-Damerell C, Aimon A, Ábrányi-Balogh P, Brandão-Neto J, Carbery A, Davison G, Dias A, Downes TD, Dunnett L, Fairhead M, Firth JD, Jones SP, Keeley A, Keserü GM, Klein HF, Martin MP, Noble MEM, O’Brien P, Powell A, Reddi RN, Skyner R, Snee M, Waring MJ, Wild C, London N, von Delft F, Walsh MA. 2020. Crystallographic and electrophilic fragment screening of the SARS-CoV-2 main protease. Nat Commun 11:5047. doi: 10.1038/s41467-020-18709-w

Drayman N, Jones KA, Azizi S-A, Froggatt HM, Tan K, Maltseva NI, Chen S, Nicolaescu V, Dvorkin S, Furlong K, Kathayat RS, Firpo MR, Mastrodomenico V, Bruce EA, Schmidt MM, Jedrzejczak R, Muñoz-Alía MÁ, Schuster B, Nair V, Botten JW, Brooke CB, Baker SC, Mounce BC, Heaton NS, Dickinson BC, Jaochimiak A, Randall G, Tay S. 2020. Drug repurposing screen identifies masitinib as a 3CLpro inhibitor that blocks replication of SARS-CoV-2 in vitro. BioRxiv Prepr Serv Biol. doi:10.1101/2020.08.31.274639

Eastman P, Swails J, Chodera JD, McGibbon RT, Zhao Y, Beauchamp KA, Wang L-P, Simmonett AC, Harrigan MP, Stern CD, Wiewiora RP, Brooks BR, Pande VS. 2017. OpenMM 7: Rapid development of high performance algorithms for molecular dynamics. PLoS Comput Biol 13:e1005659. doi:10.1371/journal.pcbi.1005659

Emsley P, Cowtan K. 2004. Coot: model-building tools for molecular graphics. Acta Crystallogr D Biol Crystallogr 60:2126–2132. doi:10.1107/S0907444904019158

Evans PR, Murshudov GN. 2013. How good are my data and what is the resolution? Acta Crystallogr D Biol Crystallogr 69:1204–1214. doi: 10.1107/S0907444913000061

Fu L, Ye F, Feng Y, Yu F, Wang Q, Wu Y, Zhao C, Sun H, Huang B, Niu P, Song H, Shi Y, Li X, Tan W, Qi J, Gao GF. 2020. Both Boceprevir and GC376 efficaciously inhibit SARS-CoV-2 by targeting its main protease. Nat Commun 11:4417. doi:10.1038/s41467-020-18233-x

Ghahremanpour MM, Tirado-Rives J, Deshmukh M, Ippolito JA, Zhang C-H, Cabeza de Vaca I, Liosi M-E, Anderson KS, Jorgensen WL. 2020. Identification of 14 Known Drugs as Inhibitors of the Main Protease of SARS-CoV-2. ACS Med Chem Lett 11:2526–2533. doi: 10.1021/acsmedchemlett.0c00521

Harvey MJ, Giupponi G, Fabritiis GD. 2009. ACEMD: Accelerating Biomolecular Dynamics in the Microsecond Time Scale. J Chem Theory Comput 5:1632–1639. doi:10.1021/ct9000685

Humphrey W, Dalke A, Schulten K. 1996. VMD: visual molecular dynamics. J Mol Graph 14:33–38, 27-28. doi: 10.1016/0263-7855(96)00018-5

Hunter JD. 2007. Matplotlib: A 2D Graphics Environment. Comput Sci Eng 9:90–95. doi:10.1109/MCSE.2007.55

Jin Z, Du X, Xu Y, Deng Y, Liu M, Zhao Y, Zhang B, Li X, Zhang L, Peng C, Duan Y, Yu J, Wang L, Yang K, Liu F, Jiang R, Yang Xinglou, You T, Liu Xiaoce, Yang Xiuna, Bai F, Liu H, Liu Xiang, Guddat LW, Xu W, Xiao G, Qin C, Shi Z, Jiang H, Rao Z, Yang H. 2020a. Structure of M(pro) from SARS-CoV-2 and discovery of its inhibitors. Nature 582:289–293. doi:10.1038/s41586-020-2223-y

Jin Z, Zhao Y, Sun Y, Zhang B, Wang H, Wu Y, Zhu Y, Zhu C, Hu T, Du X, Duan Y, Yu J, Yang Xiaobao, Yang Xiuna, Yang K, Liu X, Guddat LW, Xiao G, Zhang L, Yang H, Rao Z. 2020b. Structural basis for the inhibition of SARS-CoV-2 main protease by antineoplastic drug carmofur. Nat Struct Mol Biol 27:529–532. doi: 10.1038/s41594-020-0440-6

Jorgensen WL, Chandrasekhar J, Madura JD, Impey RW, Klein ML. 1983. Comparison of simple potential functions for simulating liquid water. J Chem Phys 79:926–935. doi: 10.1063/1.445869

Kabsch W. 2010. XDS. Acta Crystallogr D Biol Crystallogr 66:125–132. doi:10.1107/S0907444909047337

Kneller DW, Galanie S, Phillips G, O’Neill HM, Coates L, Kovalevsky A. 2020a. Malleability of the SARS-CoV-2 3CL Mpro Active-Site Cavity Facilitates Binding of Clinical Antivirals. Struct Lond Engl 1993 28:1313–1320.e3. doi: 10.1016/j.str.2020.10.007

Kneller DW, Phillips G, O’Neill HM, Jedrzejczak R, Stols L, Langan P, Joachimiak A, Coates L, Kovalevsky A. 2020b. Structural plasticity of SARS-CoV-2 3CL Mpro active site cavity revealed by room temperature X-ray crystallography. Nat Commun 11:3202. doi:10.1038/s41467-020-16954-7

Kneller DW, Phillips G, Weiss KL, Pant S, Zhang Q, O’Neill HM, Coates L, Kovalevsky A. 2020c. Unusual zwitterionic catalytic site of SARS–CoV-2 main protease revealed by neutron crystallography. J Biol Chem 295:17365–17373. doi:10.1074/jbc.AC120.016154

Lee J, Worrall LJ, Vuckovic M, Rosell FI, Gentile F, Ton A-T, Caveney NA, Ban F, Cherkasov A, Paetzel M, Strynadka NCJ. 2020. Crystallographic structure of wild-type SARS-CoV-2 main protease acyl-enzyme intermediate with physiological C-terminal autoprocessing site. Nat Commun 11:5877. doi:10.1038/s41467-020-19662-4

Liebschner D, Afonine PV, Baker ML, Bunkóczi G, Chen VB, Croll TI, Hintze B, Hung L-W, Jain S, McCoy AJ, Moriarty NW, Oeffner RD, Poon BK, Prisant MG, Read RJ, Richardson JS, Richardson DC, Sammito MD, Sobolev OV, Stockwell DH, Terwilliger TC, Urzhumtsev AG, Videau LL, Williams CJ, Adams PD. 2019. Macromolecular structure determination using X-rays, neutrons and electrons: recent developments in *Phenix*. Acta Crystallogr Sect Struct Biol 75:861–877. doi:10.1107/S2059798319011471

Ma C, Sacco MD, Hurst B, Townsend JA, Hu Y, Szeto T, Zhang X, Tarbet B, Marty MT, Chen Y, Wang J. 2020. Boceprevir, GC-376, and calpain inhibitors II, XII inhibit SARS-CoV-2 viral replication by targeting the viral main protease. Cell Res 1–15. doi:10.1038/s41422-020-0356-z

Maier JA, Martinez C, Kasavajhala K, Wickstrom L, Hauser KE, Simmerling C. 2015. ff14SB: Improving the Accuracy of Protein Side Chain and Backbone Parameters from ff99SB. J Chem Theory Comput 11:3696–3713. doi:10.1021/acs.jctc.5b00255

McCoy AJ, Grosse-Kunstleve RW, Adams PD, Winn MD, Storoni LC, Read RJ. 2007. Phaser crystallographic software. J Appl Crystallogr 40:658–674. doi:10.1107/S0021889807021206

Snijder EJ, Decroly E, Ziebuhr J. 2016. Chapter Three - The Nonstructural Proteins Directing Coronavirus RNA Synthesis and Processing In: Ziebuhr John, editor. Advances in Virus Research, Coronaviruses. Academic Press. pp. 59–126. doi:10.1016/bs.aivir.2016.08.008

Verschueren KHG, Pumpor K, Anemüller S, Chen S, Mesters JR, Hilgenfeld R. 2008. A structural view of the inactivation of the SARS coronavirus main proteinase by benzotriazole esters. Chem Biol 15:597–606. doi: 10.1016/j.chembiol.2008.04.011

Wu F, Zhao S, Yu B, Chen Y-M, Wang W, Song Z-G, Hu Y, Tao Z-W, Tian J-H, Pei Y-Y, Yuan M-L, Zhang Y-L, Dai F-H, Liu Y, Wang Q-M, Zheng J-J, Xu L, Holmes EC, Zhang Y-Z. 2020. A new coronavirus associated with human respiratory disease in China. Nature 579:265–269. doi:10.1038/s41586-020-2008-3

Xue X, Yu H, Yang H, Xue F, Wu Z, Shen W, Li J, Bartlam M, Rao Z. 2008. Structures of two coronavirus main proteases: implications for substrate binding and antiviral drug design. J Virol 82–2515.

Yang H, Yang M, Ding Y, Liu Y, Lou Z, Zhou Z, Sun L, Mo L, Ye S, Pang H, Gao GF, Anand K, Bartlam M, Hilgenfeld R, Rao Z. 2003. The crystal structures of severe acute respiratory syndrome virus main protease and its complex with an inhibitor. Proc Natl Acad Sci U S A 100:13190–13195. doi:10.1073/pnas.1835675100

Zhang L, Lin D, Sun X, Curth U, Drosten C, Sauerhering L, Becker S, Rox K, Hilgenfeld R. 2020. Crystal structure of SARS-CoV-2 main protease provides a basis for design of improved α-ketoamide inhibitors. Science 368:409–412. doi:10.1126/science.abb3405

